# HYPERsol: flash-frozen results from archival FFPE tissue for clinical proteomics

**DOI:** 10.1101/632315

**Authors:** Dylan M. Marchione, Ilyana Ilieva, Benjamin A. Garcia, Darryl J. Pappin, John P. Wilson, John B. Wojcik

## Abstract

Massive formalin-fixed, paraffin-embedded (FFPE) tissue archives exist worldwide, representing a potential gold mine for clinical proteomics research. However, current protocols for FFPE proteomics lack standardization, efficiency, reproducibility, and scalability. Here we present High-Yield Protein Extraction and Recovery by direct SOLubilization (HYPERsol), an optimized workflow using adaptive-focused acoustics (AFA) ultrasonication and S-Trap sample processing that enables proteome coverage and quantification from FFPE samples comparable to that achieved from flash-frozen tissue (average R = 0.936).

## Body

Formalin fixation and paraffin embedding (FFPE) is a tissue preparation method common in experimental research and medicine. It is standard in all pathology departments where pathological diagnosis is based on tissue section staining and immunohistochemistry on FFPE slides. The method is over one hundred years old and yields biologically inactive samples that are stable at room temperature for decades and longer^1–3^. The ubiquity of this practice in pathology combined with the unique stability of FFPE samples has resulted in massive numbers of specimens housed in countless historical tissue archives around the world. These collections represent an invaluable treasure-trove for retrospective research and translational studies, especially when specimens are paired with medical records describing the diagnosis and course of disease. However, despite this huge potential, proteomic analysis of FFPE samples has yet to become mainstream^4^. Multiple disparate protocols for proteomic analysis of FFPE material exist and generally entail a laborious deparaffinization process requiring multiple changes of toxic solvents. In recent years alternative methods have been developed, such as the SP3-CTP method, which involves a single round of xylene-ethanol incubations before resuspension in buffer containing SDS^5^, but there has yet to be a systematic comparison of FFPE extraction methods against matched frozen tissue, and therefore the optimal method for sample preparation remains unknown^6^.

Here we present High-Yield Protein Extraction and Recovery by direct SOLubilization (HYPERsol), a standardized workflow for FFPE proteomics which combines AFA™ ultrasonication and S-Trap™ ^7^ sample processing to permit highly similar peptide and protein identifications to those achieved from paired flash-frozen tissue (“F”). We developed HYPERsol by optimizing the techniques of deparaffinization, protein solubilization, and sample preparation to peptides for proteomics analysis. We compared the standard xylene-ethanol deparaffinization procedure (“X”) to a procedure in which FFPE cores were directly solubilized in buffer containing 5% SDS (“D”). Protein extraction was performed with either probe sonication (“P”) or Covaris AFA ultrasonication (“A”). Protein was recovered and processed for liquid chromatography-mass spectrometry with either Wessel-Flügge methanol-chloroform precipitation^8^ and in-solution digestion (“M”) or with S-Traps (“S”) [Fig 1a,b].

**Figure 1.**
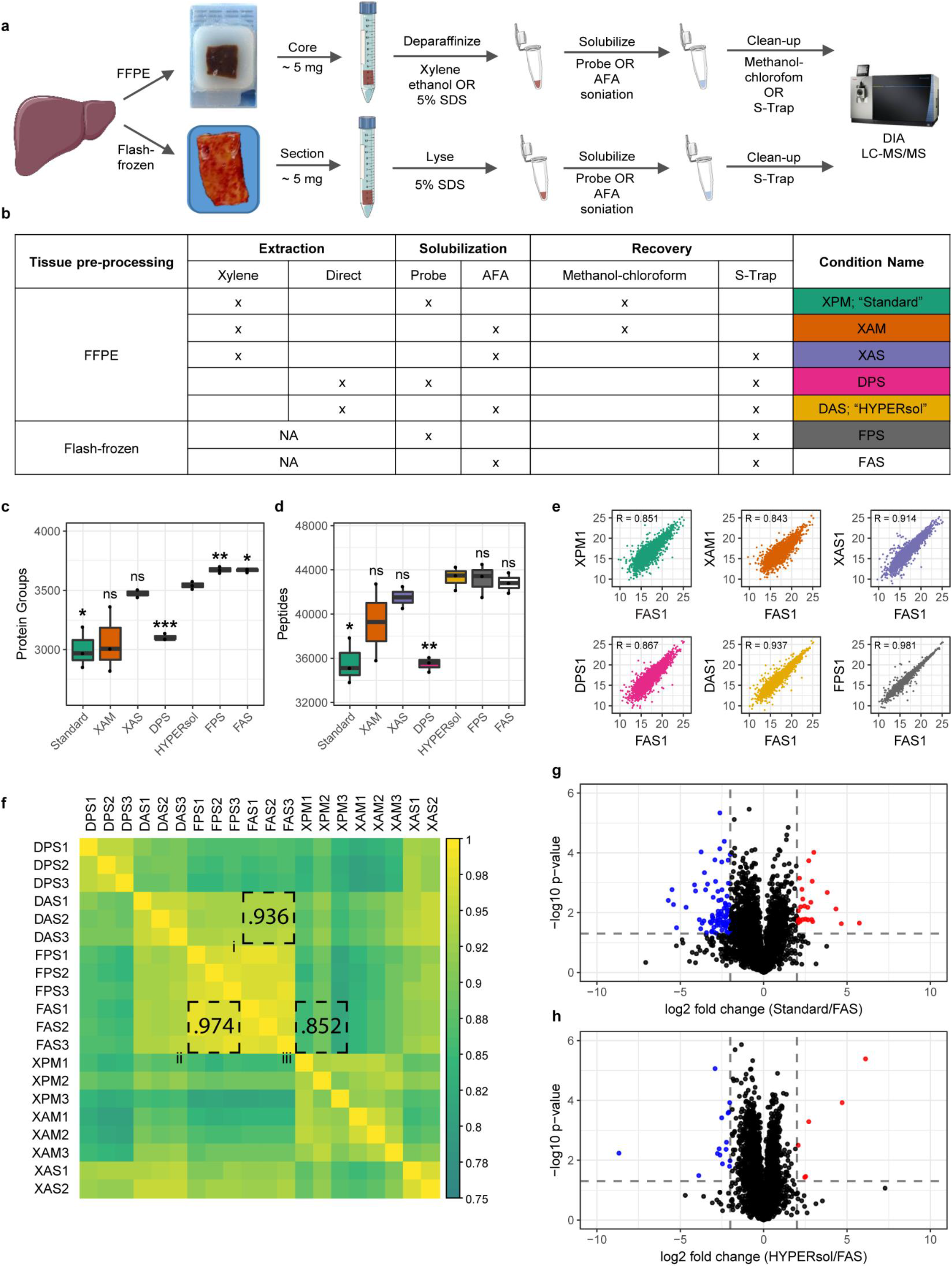
FFPE samples extracted with HYPER-sol closely resemble matched frozen tissue. **a.** Experimental workflow illustrating conditions tested. **b.** Table of experimental conditions. **c.** Tukey boxplot displaying the number of protein group IDs. **d.** Tukey boxplot displaying the number of peptide IDs. **e.** Representative scatter plots depicting the correlations between normalized protein intensity values of one replicate of each workflow against one replicate of FAS. **f.** Correlation matrix based on normalized protein intensity tables. Box i indicates the comparisons between HYPERsol and FAS. Box ii indicates the comparisons between FAS against FPS. Box iii indicates comparisons between Standard and FAS. **g.** Volcano plot comparing the relative abundance of proteins detected in Standard and FAS. **h.** Volcano plot comparing the relative abundance of proteins detected in HYPERsol and FAS. For box plots, (n = 5) asterisks indicate statistical significance compared to HYPERsol with Welch’s two-tailed t-test and p < 0.05 = *, p < 0.01 = **, p < 0.001 = ***. For volcano plots, dotted lines indicate significance thresholds with absolute log2 fold change > 2, p < 0.05 (Welch’s two-tailed t-test).

We first examined the extent of solubilization of FFPE liver samples achieved by each method. Compared to the standard workflow employing xylene-ethanol deparaffinization and probe sonication (“XP”)^9^, direct resuspension in 5% SDS buffer followed by AFA sonication (“DA”) solubilized >2-fold more protein with a corresponding >2-fold decrease in residual insoluble material [Supplementary Figure 1a-c]. To further benchmark the improved protein solubilization method we tested its ability to solubilize cores from 18 different FFPE human tissue samples. The method solubilized between 64% - 96% of the samples by mass, and the yield of soluble protein ranged from 40 - 116 μg per mg of dry FFPE, suggesting that this protocol may be compatible with workflows that require significant amounts of starting material such as post-translational modification enrichments (e.g. acetyl- or phosphoproteomics) [Supplementary Figure 1d,e].

To directly compare the performance of each sample preparation workflow for proteomic analysis against a standard tissue source, we utilized tissue from five human livers. Each sample was split at the time of autopsy: a portion was immediately flash-frozen, and another portion fixed with formalin, processed and embedded according to standard histopathology protocols. For proteomic analysis, samples were run on a Thermo Easy nLC coupled to a Thermo Fusion Tribrid mass spectrometer with a 90-minute data-independent acquisition (DIA) method. Database searching was performed in Spectronaut against a custom spectral library generated by data-dependent acquisition (DDA) analysis of a pooled sample set subject to high pH reverse phase fractionation.

On average, compared to the xylene-ethanol deparaffinization, probe sonication and methanol-chloroform precipitation (“XPM”; “Standard”), the combination of direct 5% SDS solubilization, AFA ultrasonication, and S-Trap sample processing (“DAS”; “HYPERsol“) resulted in the identification of 37% more peptides (from 30432 ± 1324 to 41643 ± 1012) which and 24% more protein groups (from 2653 ± 87 to 3297 ± 46), a depth closely approaching that obtained in a flash-frozen sample processed with probe sonication and S-Traps (FPS; 3517 ± 18) [Supplementary Figure 2a-c; Supplementary Table 1]. Among the Standard, HYPERsol, and FPS datasets the protein group overlap was 84.1%, and an additional 11% of proteins were identified in HYPERsol and FPS, but not Standard [Supplementary Figure 2d]. The average protein sequence coverage ranged from 20.0% to 23.9%, with both DPS and HYPERsol enabling statistically significant increases compared to the Standard workflow [Supplementary Figure 2e]. The distribution of gene ontology (GO) component terms was similar across all sample preparation methods, but subtle statistically significant differences were observed when comparing FFPE conditions to FPS. Extracellular, cytoplasmic, and membrane proteins were modestly overrepresented in FFPE samples relative to FPS. These differences were minimized by the HYPERsol method [Supplementary Figure 2g,h, Supplementary Table 2].

Counterintuitively, both the HYPERsol and direct solubilization, probe sonication, S-Trap (“DPS”) workflows yielded slightly more peptide identifications (IDs) than the FPS workflow [Supplementary Figure 2b]. In addition, there were 4567 peptides identified in both Standard and HYPERsol, but not FPS [Supplementary Figure 3a]. Further inspection revealed these “extra” peptides to be primarily the result of missed tryptic cleavages of highly abundant proteins, an effect presumably resulting from formalin crosslinking blocking trypsin access. The distribution of grand average of hydropathy (GRAVY) scores across the two sample sets was indistinguishable [Supplementary Figure 3b-d; Supplementary Table 3]. When lysine monomethylation and methylolation, two known chemical artifacts of formalin fixation, were included as variable modifications during both spectral library generation and searching, they were observed to be enriched in all FFPE samples, affecting approximately 5% and 0.15% of all detected peptides, respectively [Supplementary Figure 3e,f, Supplementary Table 3]^10^. Like missed cleavages, these modifications were primarily detected on highly abundant proteins, and omitting them from database searching did not greatly alter the library composition [Supplementary Figure 3g,h]. All together, these results suggest that, despite the persistent presence of artifacts induced by formalin, HYPERsol sample processing enables deeper and more reproducible FFPE proteome analysis than the former standard workflow.

Next, in order to evaluate the extent to which protein extracted from FFPE samples resembles matched frozen tissue, we directly compared each method against multiple liver samples derived from a single patient. We introduced two additional conditions: “XAS” (xylene-ethanol, AFA, S-Trap) to determine if the combination of AFA and S-Trap processing could improve the data quality from xylene-deparaffinized samples, and “FAS” (flash-frozen, AFA, S-Trap) to determine if AFA sonication would enable deeper proteome coverage than probe sonication on flash-frozen tissue. Samples were analyzed on a Thermo Dionex Ultimate 3000 coupled to a Thermo QE HF-X, and the DIA method was extended by 20 minutes to enable slightly deeper coverage. Methyl and methylol-lysine adducts were not included in the search. Again, among FFPE conditions, the HYPERsol protocol resulted in the greatest number of peptide and protein group IDs [Figure 1c,d; Supplementary Table 4]. The numbers of peptides and protein groups identified in the XAS samples were markedly better than those from the former standard workflow, indicating that the combination of AFA sonication and S-Trap cleanup can partially compensate for the inefficiency of xylene-ethanol deparaffinization. Surprisingly, substitution of the probe sonication with AFA did not improve the number of peptide or protein group IDs from the flash-frozen tissue, suggesting that the increased sonication energy afforded by AFA is only obligatory when solubilizing protein from FFPE. Nevertheless, given the improved performance of AFA relative to probe sonication in the other conditions, we considered the FAS dataset as the ground truth dataset for subsequent comparisons.

In order to compare the similarity of the proteomic datasets generated via each sample preparation workflow, we examined the Pearson correlation coefficients among the protein quantification tables derived from each workflow [Supplementary Table 5, Figure 1e,f] against FAS. Overall, the HYPERsol datasets were exceptionally similar to the FAS datasets, with R ranging from 0.928 to 0.951 across any pairwise comparison, and an average R of 0.936 [Figure 1f, box i]. The average correlation between the FAS and FPS datasets was 0.974, indicating that the variability introduced by FFPE and HYPERsol extraction is not substantially greater than that introduced by alternative sample preparation strategies for replicates of the same flash-frozen tissue [Figure 1f, box ii]. Analyses of protein extracted with the standard method were less similar (range = 0.811 - 0.890, average = 0.852), suggesting that in addition to reducing the overall number of IDs, the incomplete solubilization and recovery afforded by the former standard workflow also distorts the relative abundance of the detected proteins [Figure 1f, box iii]. To further explore this possibility we generated volcano plots comparing the relative abundance of protein groups that were identified across both standard and FAS [Figure 1g], or both HYPERsol and FAS extractions [Figure 1h, Supplementary Table 6]. Whereas use of the standard protocol resulted in 120 proteins with absolute log_2_ fold-change threshold > 2 and p-val < 0.05 out of the 2956 quantified (101 underestimated, 19 overestimated; representing 4.0% of total proteins), the use of HYPERsol reduced this number to only 24 out of 3517, (6 overestimated, 18 underestimated; representing (0.68% of total proteins) reducing experimental noise and thereby effecting a more faithful representation of the composition of the original tissue.

While many potential applications exist, proteomic analysis of FFPE tissue is especially well-suited to identify new immunohistochemical (IHC) markers to facilitate the diagnosis of tumors for which histomorphology is insufficiently specific. Malignant peripheral nerve sheath tumor (MPNST) is one such tumor which is notoriously difficult to diagnose^11^. There are no reliable positive IHC markers for MPNST. The best-established IHC targets are H3 K27 di- or trimethylation, but these marks are only globally altered in approximately half of cases^12,13^. It is particularly difficult to distinguish MPNST from histologic mimics including melanoma and synovial sarcoma [Figure 2a]^14^.

**Figure 2.**
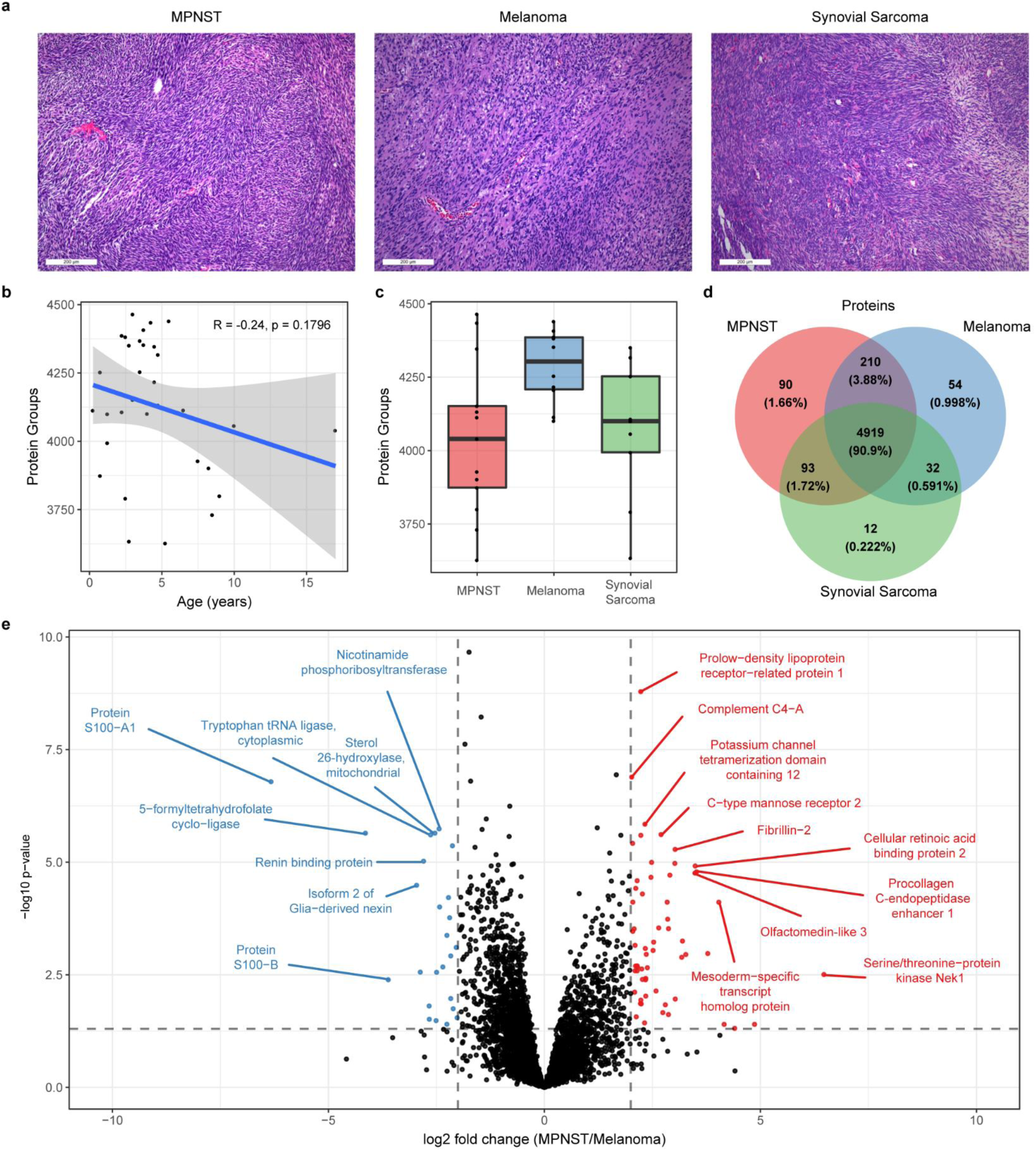
HYPER-sol enables analysis of archival FFPE tumor samples. **a.** Representative H&E stains of MPNST, melanoma, and synovial sarcoma illustrating their histomorphologic similarity. **b.** Scatterplot depicting the correlation between specimen age and the number of protein groups identified in a single 135 minute DIA run. R and p-value are obtained from Pearson’s product-moment correlation **c.** Tukey boxplot of the average number of protein group IDs from each tumor type (n = 13 MPNST, 10 melanoma, 9 synovial sarcoma). **d.** Venn diagram of protein IDs from each tumor type. **e.** Volcano plot comparing MPNST to melanoma. 4851 proteins are depicted. Dotted lines indicate significance thresholds at absolute log2 fold change > 2 and p < 0.05 (Student’s two-tailed t-test).

We therefore applied HYPERsol to 32 archival FFPE tumor samples (13 MPNSTs, 10 melanomas, and nine synovial sarcomas) to identify new candidate IHC markers. Approximately 4000 - 4300 protein groups were detected per sample [Figure 2a,b]. The oldest sample had been in storage for over 17 years, and there was no correlation between the duration of storage and the number of protein identifications [Figure 2b; Supplementary Table 7]. Among the 5410 unique protein groups identified overall, approximately 91% were detected in at least one case of each tumor type, a figure which numerically illustrates the inherent difficulty of identifying distinguishing markers [Figure 2d; Supplementary Table 8].

Nevertheless, volcano plots revealed proteins that were highly enriched in each tumor type, and these were largely consistent with known protein expression signatures [Figure 2e, Supplementary Figure 4a,b, Supplementary Table 9]. For example, S100A and S100B were highly overexpressed in melanomas relative to both MPNSTs and synovial sarcomas, consistent with the longstanding use of S100 as an IHC marker for melanoma, and the loss of S100 expression seen in MPNST as compared to benign nerve sheath tumors [Figure 2e, Supplementary Figure 4b]^15,16^ Additionally, TLE1, a widely-used marker for synovial sarcoma, was indeed detected in all synovial sarcomas^17^. However, in line with reports that TLE1 is not entirely specific for synovial sarcoma, TLE1 expression was also observed in 9/10 melanomas and 12/13 MPNSTs^18^. Though statistically significant, on average it was only approximately 3-fold more highly expressed in synovial sarcoma relative to MPNST, an expression difference that further underscores the need for better markers to distinguish between these two tumors [Supplementary Figure 4a]. Importantly, this experiment revealed several proteins that were entirely tumor-type specific (90 in MPNST, 54 in melanoma, and 12 in synovial sarcoma; [Supplementary Table 9]). Future work will establish whether these are useful IHC markers for these diagnostically challenging tumors.

In conclusion, HYPERsol enables highly reproducible protein identification and quantification from FFPE tissue, yielding results that are highly similar to flash-frozen tissue. The combination of direct SDS solubilization, AFA ultrasonication and S-Trap sample processing markedly increases protein yield, potentially enabling analysis of small samples such as core needle biopsies or samples obtained by laser-capture microdissection. We anticipate that the reduced variability of HYPERsol sample processing will enhance the capacity of researchers to extract meaningful biologic information from FFPE samples by minimizing sample preparation-induced artifacts. In addition, the approach is rapid and obviates the need for time-consuming, tedious and toxic deparaffinization: samples can now be easily prepared to peptides and analyzed on the same day. With the availability of 96-well plates for AFA and S-Trap sample processing, HYPERsol is suited to automated, high-throughput analyses essential for clinical implementation. We thus anticipate that the HYPERsol workflow will enable novel discoveries from rich clinically-annotated and histologically-characterized FFPE biorepositories worldwide, thereby helping to usher in a new era of clinical proteomics.

## Methods

### IRB statement

Human samples were collected under protocols approved by the Institutional Review Board (IRB) of the University of Pennsylvania or documented as exempt from IRB review. All samples were subjected to histopathologic review for confirmation of diagnosis and selection of the region for tissue isolation.

### Tissue processing

Automated tissue processing was carried out in a Leica Peloris II processor (Leica Biosystems) with the following incubation settings: 60 min 10% neutral buffered formalin, 60 min formalin 10% neutral buffered formalin, 80% EtOH 20 minutes, 95% EtOH 60 minutes, 100% EtOH 30 minutes, 100% EtOH 50 minutes, 100% EtOH 60 minutes, xylene 30 minutes, xylene 50 minutes, xylene 60 minutes, paraffin 60 minutes, paraffin 60 minutes, paraffin 60 minutes. Following processing, samples were embedded in paraffin and stored in blocks at room temperature prior to processing.

### Hematoxylin & Eosin staining

The original diagnostic histologic sections were used for confirmation of diagnosis and for the photomicrographs shown in Figure 2. These were standard 5 μm tissue sections stained with hematoxylin/eosin according to standard histopathology protocols. Digital images were taken on a Leica DMC 4500 camera and captured and processed using the Leica Digital Application Suite v4.12.

### Deparaffinization/solubilization

#### Xylene/ethanol

Tissue cores were diced into small pieces and resuspended in 10 x volume (based on dry weight) of xylene and incubated at 37 °C for 10 minutes with gentle agitation. Following centrifugation and removal of xylene, the process was repeated with xylene and then with 100% ethanol twice, 95% ethanol, 85%, 70%, 50%, 20%, and then water. Samples were then resuspended in solubilization buffer containing 5% SDS and 100 mM Tris pH 8.5, and homogenized with a mortar and pestle. Samples were then passed through an 18-gauge needle 10x, then a 21-gauge needle 10x. Following sonication (as described below), the process was repeated. The homogenized lysate was then spun down at 16,000 x g in a benchtop centrifuge for 15 minutes. The soluble fraction was transferred to a separate tube and the total protein concentration was measured using a BCA assay (Pierce).

#### Direct solubilization

Cores were resuspended in 20x volume/weight of solubilization buffer containing 5% SDS and 100 mM Tris pH 8.5, and incubated at 50 °C for 10 min. The pellet was homogenized with a micropestle, then passed through an 18-gauge needle 10x, followed by a 21-gauge needle 10x. Following sonication as described below, the samples were placed on a heat block at 80 °C for 1 hr. The samples were removed and the sonication was repeated. The samples were returned to the heating block for 1 hr. The homogenized lysate was then spun down at 16,000 x g in a benchtop centrifuge for 15 minutes. The soluble fraction was transferred to a separate tube and the total protein concentration was measured using a BCA assay (Pierce).

### Sonication

#### Probe

For probe sonication, the homogenized lysate was subjected to benchtop sonication with a Thermo Fisher probe sonicator with a microtip, 3 × 30 s pulses, 20% power, and with a 50% duty ratio. Two rounds of sonication were performed as described above.

#### AFA

Samples were analyzed on a Covaris S220 AFA in screw-cap microTUBEs (PN500339). The general parameters were as follows: water level set point 15, chiller set point 18 °C, holder, peak incident power 175 W, duty factor 10%, cycles per burst 200, instrument temperature 20 °C. Two rounds of sonication were performed; in the first round (emulsification), the treatment time was 300 s. In the second round (solubilization), the treatment time was 360 s.

### Tissue yield

To calculate tissue yield and percent solubilized for each tissue type depicted in Figure S2, dried 1 mm tissue cores were weighed, then directly placed in solubilization buffer, ground with a mortar and pestle, and allowed to equilibrate for 15 minutes at room temperature. The sample was spun at 16,000 x g in a benchtop centrifuge and the initial supernatant was saved for subsequent BCA assay. The pellets were also weighed and recorded as the resolubilized mass of the FFPE tissue. The samples were then processed according to the HYPERsol protocol, and spun down again at 16,000 x g in a benchtop centrifuge. The supernatant was saved for BCA assay, and the pellet re-weighed. The % solubilized was calculated as: 100*(1-((mass of residual pellet after processing)/(mass of starting pellet after equilibration in DAS/HYPERsol buffer))). The yield was calculated as: (protein concentration × volume for the initial resuspension solution + protein concentration x volume for the final, processed sample) / (initial weight of dried FFPE in mg). For all samples, the amount of protein in the initial resuspension/equilibration solution contributed negligibly to the overall yield.

### Sample clean-up and preparation for LC-MS

#### Methanol/chloroform

Samples were processed as previously described^8^. The sample volume containing 100 μg of total protein was adjusted to 300 μl with water. 300 μl of ice-cold methanol was added, and the sample was vortexed briefly. Next, 75 μl of ice-cold chloroform was added and the sample was vortexed again, then immediately centrifuged for 1 min at 9,000 x g on a benchtop centrifuge. Following centrifugation, the upper phase was removed and discarded. 300 μl of ice cold methanol was added and the sample was vortexed again, then spun at 16,000 x g in a benchtop centrifuge. The supernatant was removed without disturbing the pellet, and the tube was left with the cap off for 5 minutes to allow excess methanol to evaporate. The pellet was then resuspended in 20 μl of 6 M urea + 2 M thiourea. Following resuspension, 100 μl of 50 mM ammonium bicarbonate pH 8.0 with 1.2x protease inhibitor cocktail (Pierce) was added, and DTT was added to a final concentration of 10 mM. Samples were incubated for 30 minutes at room temperature. Iodoacetamide was added to a final concentration of 20 mM and samples were incubated in the dark for 30 minutes. An additional 10 mM DTT was added to quench the derivatization reaction, and the samples were digested with trypsin at a 1:50 ratio overnight at room temperature.

#### S-Trap

Prior to loading on the S-Traps, samples were reduced and alkylated as described above. S-Trap trypsin digestion and clean-up was performed according to the manufacturer’s instructions. Briefly, 50 μg protein was loaded on S-Trap micro spin columns (C02-micro) and washed extensively with 90% methanol, 100 mM TEAB, pH 7.1. Trypsin was added directly to the microcolumn at a 1:20 ratio in 50 mM TEAB, pH 8, and samples were incubated in a water bath at 47 °C for 1 hr. Peptides were eluted by serial addition of 50 mM TEAB, 0.2% formic acid, and 0.2% formic acid in 50% acetonitrile.

#### High-pH reversed-phase fractionation

A fraction of the peptides from each sample were pooled and acidified with 0.1% trifluoroacetic acid (TFA). The pooled mix was loaded on a Harvard apparatus Micro SpinColumn (Cat# 74-4601), washed with 0.1% TFA, and eluted with 12 serial additions of 100 mM ammonium formate, pH 10, containing increasing concentrations of acetonitrile (10, 12, 14, 16, 18, 20, 22, 24, 26, 28, 35, 60% ACN). Fractions were pooled (1+6, 2+7, etc.), dried, and desalted prior to analysis.

#### Desalting

All samples were resuspended in 0.1% TFA, loaded on homemade C18 stage-tips (3M Empore Discs) and desalted as previously described, with minor modifications^19^. Briefly, columns were conditioned with 100 μl acetonitrile and equilibrated with 100 μl 0.1% TFA. Samples were loaded and the stage-tip was washed with 100 μl 0.1% TFA before peptides were eluted with 100 μl 0.1% formic acid in 60% acetonitrile.

### Liquid chromatography

Nanoflow liquid chromatography was performed using either a Thermo Dionex Ultimate 3000 or a Thermo ScientificTM Easy nLC™ 1000 equipped with a 75 µm x 20 cm column packed in-house using Reprosil-Pur C18-AQ (2.4 µm; Dr. Maisch GmbH, Germany). Buffer A was 0.1% formic acid and Buffer B was 0.1% formic acid in 80% acetonitrile. The flow rate was 400 nL/min.

For 90 minute runs (Easy nLC) the gradient was as follows: 2% B for 2 minutes, 18% B over 42 minutes, 40% B over 30 minutes, 46% B over 4 minutes, 55% B over 2 min, 98% B over 10 s, hold 98% B for 9 min 50 s.

For 110 minute runs (Dionex) the gradient was as follows: 5% B for 1 minute, 30% B over 73 minutes, 45% B over 18 minutes, 85% B over 1 minute, hold for 7 minutes, re-equilibrate for 10 minutes.

For 135 minute runs (Easy nLC) the gradient was as follows: 1% B to 4% B over 3 minutes, 6% over 3 minutes, 8% over 4 minutes, 10% over 5 minutes, 12% over 18 minutes, 17% over 9 minutes, 26% over 41 minutes, 28% over 9 minutes, 30% over 6 minutes, 32% over 5 minutes, 34% over 4 minutes, 36% over 4 minutes, 38% over 3 minutes, 41% over 3 minutes, 52% over 3 minutes, 90% over 5 minutes, hold 90% for 10 minutes.

### Mass spectrometry

The HPLC was coupled online to either a Thermo Fusion Orbitrap Tribrid or a Thermo Q Exactive HF-X mass spectrometer operating in the positive mode using a Nanospray Flex™ Ion Source (Thermo Scientific) at 2.3 kV.

DDA analysis on the Fusion was performed with the following settings: MS1 350-1200 m/z, 120K resolution, AGC target 1e6, max inject time 60 ms; MS2 15K resolution, AGC 5e4, max inject time 120 ms, Top Speed = 3 s, isolation window = 2 m/z, stepped HCD collision energy 29 +/− 5%, include z = 2-5.

DIA analysis on the Fusion was performed with the following settings: MS1 350-1200 m/z, 120K resolution, AGC target 1e6, max inject time 60 ms; MS2 30K resolution, AGC 1e6, max inject time 54 ms, 33 windows of 25.7 m/z, stepped HCD collision energy 27 +/− 5%.

DDA analysis on the QE HF-X was performed with the following settings: MS1 350-1200 m/z, 120K resolution, AGC 3e6, max inject time 50 ms; MS2 30K resolution, AGC 1e5, max inject time 120 ms, Top N = 20, isolation window 1.3 m/z, stepped nce (25.5, 27, 30), exclude z = unassigned and z = 1.

DIA analysis on the QE HF-X was performed with the following settings: MS1 350-1200 m/z, 120K resolution, AGC 3e6, max inject time 50 ms; MS2 30K resolution, AGC 3e6, max inject time auto, MSX 1, 45 windows with a width of 20 m/z and margins of 0.5, stepped nce (25.5, 27, 30).

### Mass spectrometry data analysis

For all liver sample analyses, spectral libraries were generated using Spectronaut Pulsar X with the following settings: digest type = specific, missed cleavage = 2, min peptide length = 7, max peptide length = 52, toggle N-terminal M = true, and using the 2017-10-25 version of the Homo sapiens [SwissProt TaxID=9606] proteome. Liver samples were also analyzed with directDIA and the search archives were used to improve the Pulsar library search. For the methyl and methylol adduct search, these were included as variable modifications. For histologic mimics, the spectral library was generated with Proteome Discoverer 2.3 using the default “PWF_QE_SequestHT_MSAmanda_Percolator” and “CWF_Basic” workflows. The PDRESULT file was converted to a spectral library in Spectronaut Pulsar X with the settings described above.

All DIA runs were analyzed in Spectronaut Pulsar X using BGS Default Factory Settings. Peptide and protein intensities were log2 transformed, and processed by a two-step normalization. First, within each run, the run-level median intensity was subtracted from each measured intensity such that the values were normally distributed around 0. Then the global median intensity from the entire sample set was added back such that all intensity values were positive. No imputation was performed. Figures were generated in R using the packages ggplot2, ggthemes, corrplot, and VennDiagram.

## Author contributions

JBW and JPW conceived the work. JBW, JPM, DM, and DP designed the experiments. DM, JBW, and II acquired the data. DM, JBW, and JPW analyzed the data. JBW, BAG, and JPW supervised aspects of the work. DM, JBW, and JPW were primarily responsible for writing and revising the manuscript. All authors approved the final version of the manuscript.

## Supplementary Figures

**Supplementary Figure 1.**
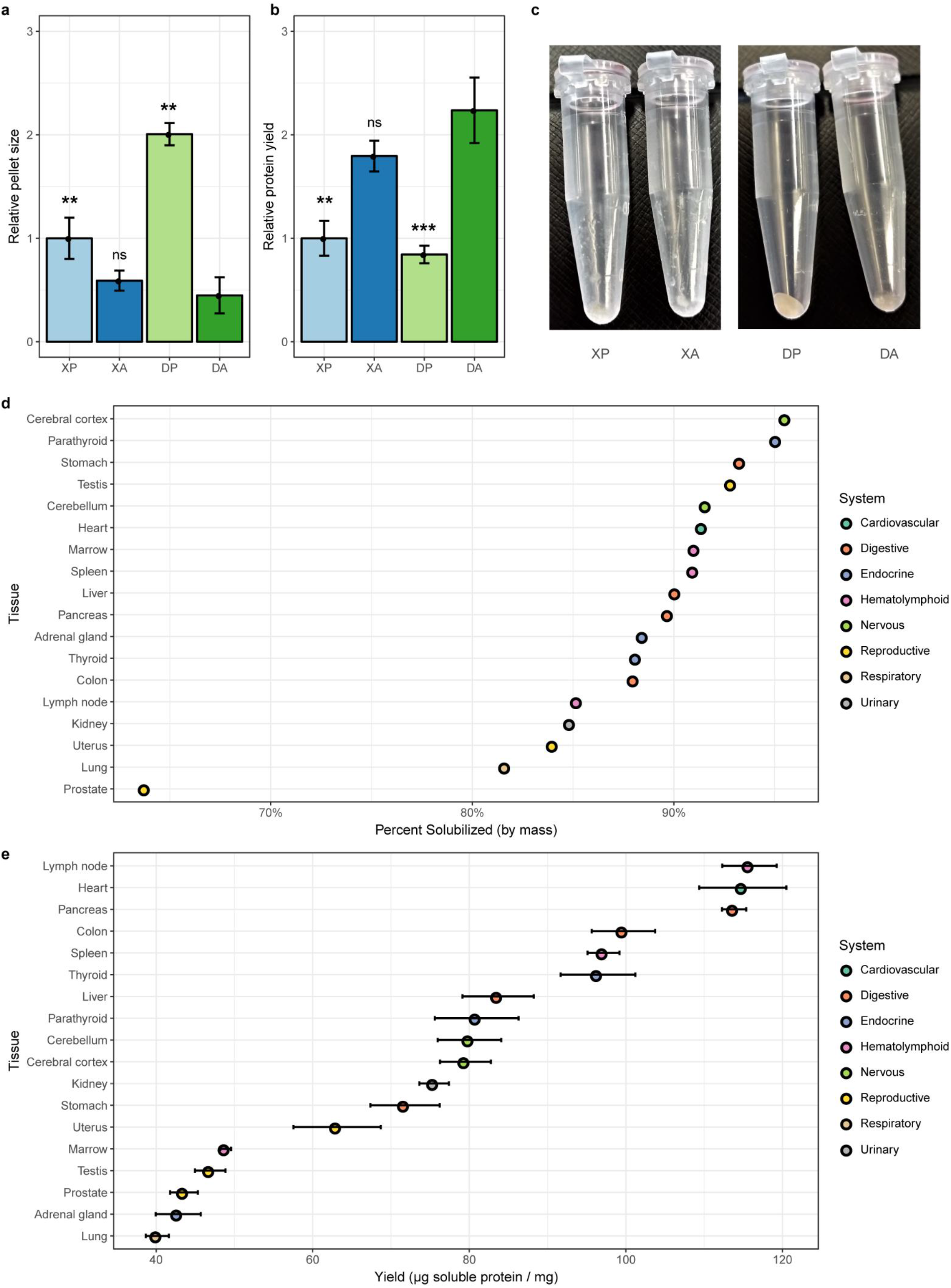
Direct 5% SDS solubilization combined with AFA sonication is an effective FFPE tissue solubilization strategy. **a.** Bar graph of relative residual pellet mass, normalized to initial pellet mass. **b.** Bar graph of relative protein yields, normalized to initial pellet mass. **c.** Representative images of residual material after each solubilization/sonication technique. **d.** Scatter plot depicting the extent of solubilization of 18 FFPE human tissue samples (n = 1). **e.** Scatter plot depicting protein yield per milligram of dry FFPE across 18 FFPE human tissue samples (n = 1). Error bars depict the standard deviation from three replicate BCA assays. For bar graphs, means ± standard deviations are shown (n = 4), and asterisks indicate statistical significance compared to HYPER-sol with Welch’s two-tailed t-test and p < 0.05 = *, p < 0.01 = **, p < 0.001 = ***.

**Supplementary Figure 2.**
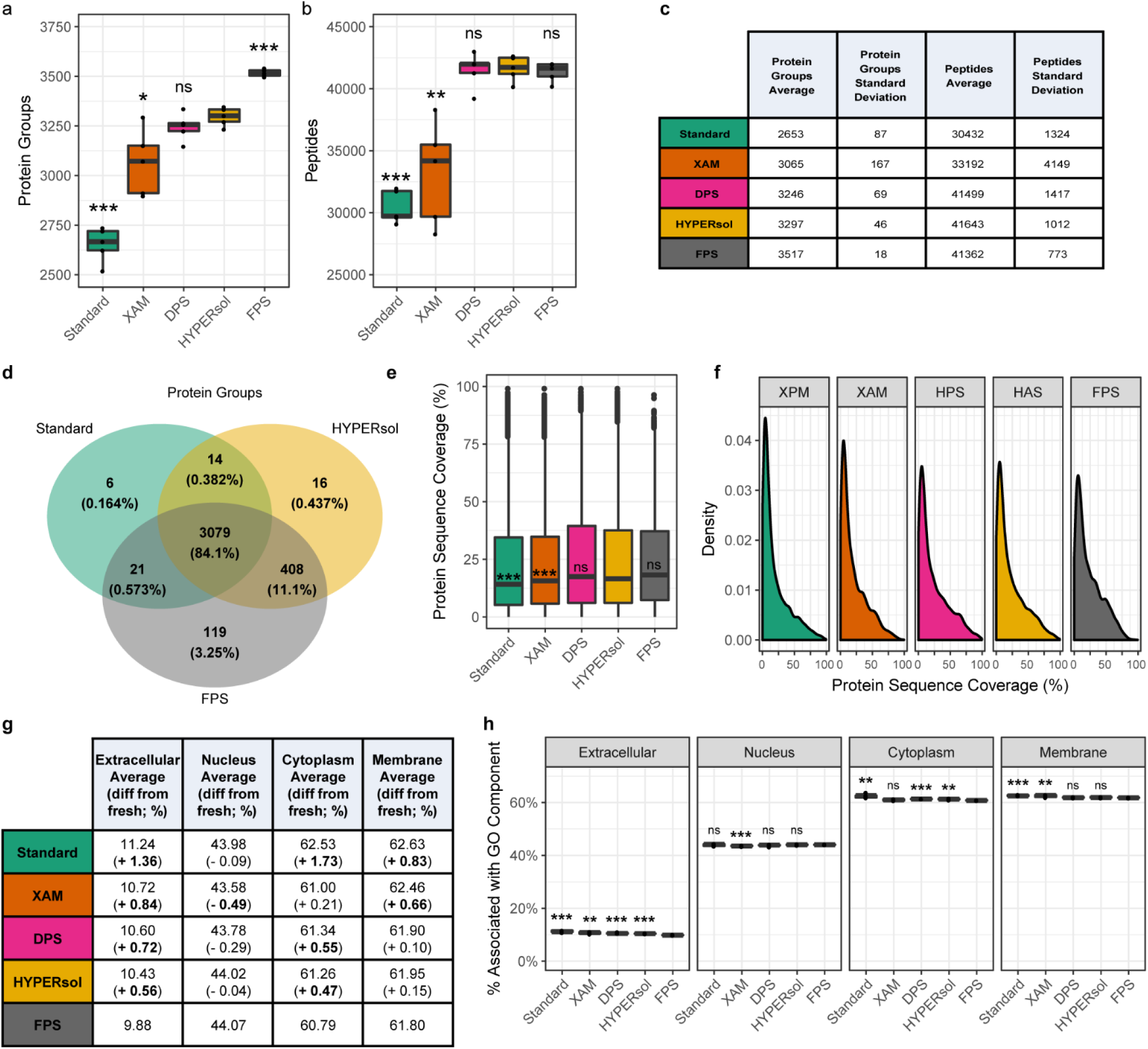
The combination of direct SDS solubilization, AFA sonication, and S-Trap sample processing (HYPERsol) enables deeper and more reproducible FFPE proteome analysis than the current standard workflow. **a.** Tukey boxplot displaying the number of protein group IDs. **b.** Tukey boxplot displaying the number of peptide IDs. **c.** Table displaying data used for panels a and b. **d.** Venn diagram displaying overlap of protein groups among Standard, HYPERsol, and FPS. **e.** Tukey boxplot displaying protein sequence coverages across each experimental condition. **f.** Density plot portraying the distribution of sequence coverages across each condition. **g.** Table depicting the fraction of identified proteins associated with each GO component term across conditions. **h.** Bar plot depicting the data from panel g. Asterisks indicate statistical significance compared to FPS with Welch’s two-tailed t-test and p < 0.05 = *, p < 0.01 = **, p < 0.001 = ***.

**Supplementary Figure 3.**
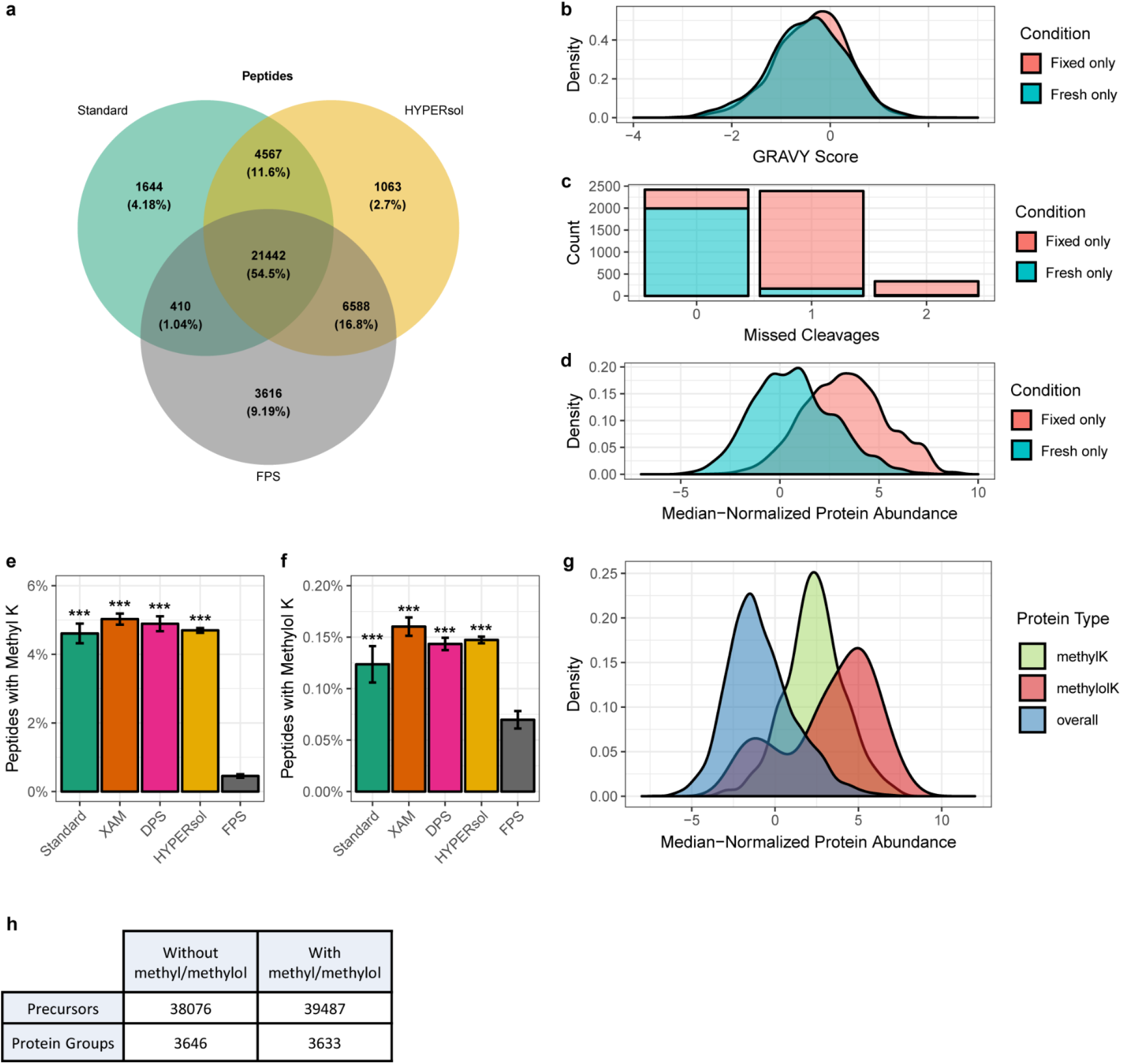
Unique peptides detected from FFPE contain missed cleavages and artefactual modifications. **a.** Venn diagram of peptide IDs comparing HYPERsol, the standard workflow, and FPS. **b.** Distribution of grand average of hydropathy (GRAVY) scores for peptides detected in all FFPE samples but never in flash-frozen samples (“Fixed only”) and peptides detected in all flash-frozen samples but never in FFPE samples (“Fresh only”).. **c.** Histogram of missed cleavage counts among condition-specific peptides **d.** Distribution of the protein abundances for condition-specific peptides. **e.** Fraction of peptides with monomethyl lysine. **f.** Fraction of peptides with methylol lysine. **g.** Distribution of the average abundance of detected proteins, stratified by modification state (overall, methylated, and methylolated). **h.** Table depicting the effect of including variable formalin-induced lysine modifications during database searching on spectral library size. For bar graphs, means ± standard deviations are shown, (n = 5) and asterisks indicate statistical significance compared to FPS with Welch’s two-tailed t-test and p < 0.05 = *, p < 0.01 = **, p < 0.001 = ***.

**Supplementary Figure 4.**
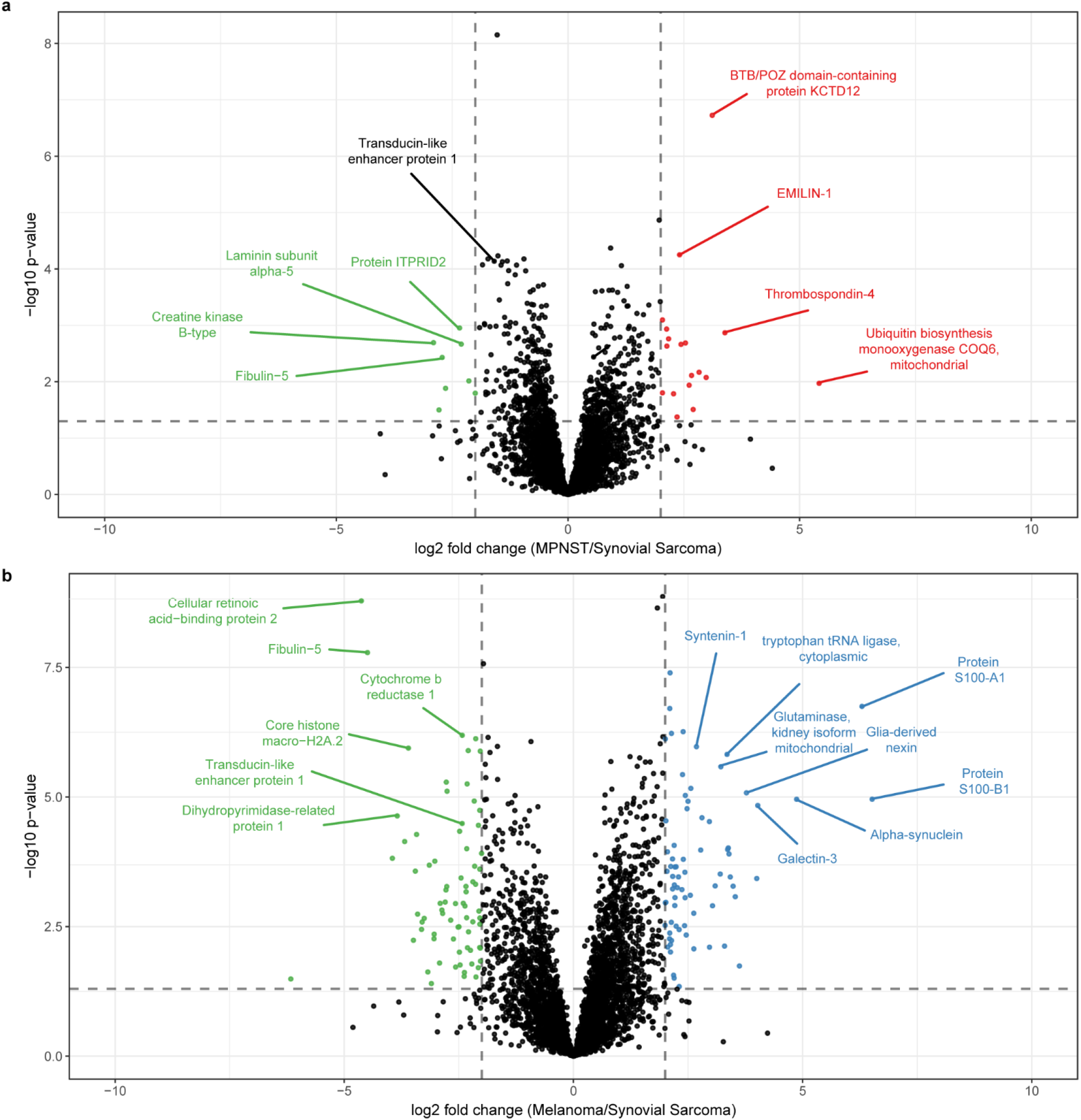
HYPER-sol enables analysis of archival FFPE tumor samples. **a.** Volcano plot comparing MPNST to synovial sarcoma. 4737 proteins are depicted. **b.** Volcano plot comparing melanoma to synovial sarcoma. 4640 proteins are depicted. For volcano plots, dotted lines indicate significance thresholds at absolute log2 fold change > 2 and p < 0.05 (Student’s two-tailed t-test).

## Supplementary Tables

Table S1. Experiment 1 Peptide and Protein Reports

Table S2. Experiment 1 Gene Ontology Terms

Table S3. Experiment 1 Unique Peptides and Protein Modifications

Table S4. Experiment 2 Peptide and Protein Reports

Table S5. Experiment 2 Protein Correlation Matrix

Table S6. Experiment 2 Volcano Plots

Table S7. Archival FFPE Tumor Specimen Age and Number of Protein IDs

Table S8. Archival FFPE Tumor Specimen Protein Report

Table S9. Archival FFPE Tumor Specimen Volcano Plots and Specific Proteins

